# Characterisation of morphological differences in well-differentiated nasal epithelial cell cultures from preterm and term infants at birth and one-year

**DOI:** 10.1101/370148

**Authors:** Helen E. Groves, Hong Guo-Parke, Lindsay Broadbent, Michael D. Shields, Ultan F. Power

## Abstract

**Background:** Innate immune responses of airway epithelium are important defences against respiratory pathogens and allergens. Newborn infants are at greater risk of severe respiratory infections compared to older infants. However, very little is known regarding human neonatal airway epithelium immune responses and whether age-related morphological and/or innate immune changes contribute to the development of airway disease.

**Methods:** We collected nasal epithelial cells from 41 newborn infants (23 term, 18 preterm) within 5 days of birth. Repeat sampling was achieved for 24 infants (13 term, 11 preterm) at a median age of 12.5 months. Morphologically and physiologically authentic well-differentiated primary paediatric nasal epithelial cell (WD-PNEC) cultures were generated and characterised using light microscopy and immunofluorescence.

**Results:** WD-PNEC cultures were established for 15/23 (65%) term and 13/18 (72%) preterm samples at birth, and 9/13 (69%) term and 8/11 (73%) preterm samples at one-year. Newborn and infant WD-PNEC cultures demonstrated extensive cilia coverage, mucous production and tight junction integrity. Newborn WD-PNECs took significantly longer to reach full differentiation and were noted to have much greater proportions of goblet cells compared to one-year repeat WD-PNECs. No differences were evident in ciliated/goblet cell proportions between term- and preterm-derived WD-PNECs at birth or one-year old.

**Conclusion:** WD-PNEC culture generation from newborn infants is feasible and represents a powerful and exciting opportunity to study differential innate immune responses in human airway epithelium very early in life.

## Introduction

The airway epithelium plays a crucial role in initiating airway innate immune response mechanisms in humans. It facilitates this initial response by providing a mechanical barrier to pathogen entry and releasing antimicrobial and inflammatory peptides in response to innate immune receptor stimulation by pathogens.(1) Certain respiratory disorders, including asthma and cystic fibrosis, are associated with altered airway epithelial cell (AEC) immune responses and impaired barrier function of the epithelium.(2,3) There is increasing evidence that asthma and other chronic respiratory disorders begin in early life and it is possible that airway innate immune responses undergo maturation in parallel with postnatal lung growth, differentiation, microbiome colonisation, and external infectious and non-infectious insults.(2,4) However, little is known regarding the development of these airway epithelial innate immune responses in early life.

In adults and children AECs can be obtained from brushings of the nasal or bronchial tracts and prior studies have demonstrated the successful use of both bronchial and nasal AEC cultures in investigating early life respiratory disorders.(5–8) However, undertaking bronchial brushings in very young infants is impractical, as acquiring samples could only be ethically conducted opportunistically when infants are intubated. As an alternative, the nasal passage provides an easily accessible source of AECs that is much less invasive. Recent publications have highlighted the potential benefit of this technique in investigating early changes in immune function in cystic fibrosis (CF) infants, in the study of the nasal transcriptome of infants, and in the innate immune responses of the airway epithelium to allergens as part of asthma pathogenesis research.(7,9,10) Mosler *et al* described the culture of nasal epithelial cells from CF infants, some of whom were as young as one-month old, but to date, Miller *et al* is the only publication describing monolayer culture of nasal epithelial cells from newborn infants.(7)

One of the major drawbacks of studying human infant primary AECs is the challenge of reduced proliferation after a small number of cell passages. As a possible strategy to deal with this challenge, Wolf *et al* described the generation of conditionally reprogrammed cells (CRC) from harvested infant AECs with enhanced proliferative and survival capacity.(11) However, generating CRCs requires alteration of primary AECs and it remains unclear what impact this might have on AEC innate immune responses.

A further development in AEC culture has been the creation of differentiated epithelial cell cultures via formation of an air-liquid interface. Indeed, we previously described the formation of well-differentiated paediatric nasal airway epithelial cell cultures (WD-PNECs) from older infants and demonstrated this model reproduces many of the hallmarks of respiratory syncytial virus (RSV) cytopathogenesis seen *in vivo*.(6)

Young infants, especially those born prematurely, have increased susceptibility to severe respiratory disease following infections, such as respiratory syncytial virus infection (RSV).(12,13) Neonates in particular have much greater susceptibility to a variety of infections compared to older infants and adults.(14) Furthermore, encounters with pathogens during the crucial period of neonatal development may have long term impacts on future respiratory health.(15) For instance, it is known that severe RSV infection during early infancy is linked to later diagnosis of wheeze and asthma(16,17) and even later life respiratory diseases, such as chronic obstructive pulmonary diseases (COPD), may be associated with early life experiences.(4) Therefore, given the current limited knowledge of the development of early AEC immune responses, greater understanding of these responses is needed to yield novel insights into the mechanisms of susceptibility underpinning childhood airway disease.

Here we report the first successful generation of WD-PNECs from infants at birth and compare this with WD-PNECs derived from the same infants at one-year old. This paper details the characterisation of this novel newborn airway model and describes the morphological differences between newborn and older infant airway epithelial cultures. Our work, therefore, presents an exciting opportunity to study ‘‘naive’’ human airway epithelial cells in early life and to investigate the developmental immunobiology of the airway epithelium over the first year of life.

## Methods

### Subjects and study design

Healthy newborn term infants, (37-42 weeks gestation) and preterm infants (28-34 weeks gestation) underwent a nasal brushing procedure up to 5 days after birth (median age 2 days, range 6 hours to 5 days) at the Royal Jubilee Maternity Hospital, Belfast. Infants with known severe congenital anomaly of the airway, immunodeficiency, or congenital heart disease at the time of recruitment were excluded. A repeat nasal brushing sample was taken from a subset of infants at one-year old and a medical questionnaire recording previous episodes of upper/lower respiratory tract symptoms and/or bronchiolitis was completed based on parent recall (see supporting information, S1 Appendix).

### Sampling of nasal epithelial cells

Nasal airway epithelial cells (AECs) were harvested from healthy, non-sedated neonates at the earliest opportunity post-delivery. Nasal sampling was performed either with the neonate lying in a parent’s arms or in a cot using the technique described by Miller *et al*.(7) In brief, the infant’s head was gently secured using one hand and a 2.7 mm diameter interdental brush (DentoCare Professional, London, UK) was introduced into each nostril (one brush/nostril) in turn and gently rotated twice against the medial aspect of the inferior turbinate.

We collected nasal AECs from 41 newborn infants (23 term, 18 preterm) within the first 5 days of life (median age 2 days, range 6 hours to 5 days) (Table 1). The procedure was well tolerated by all neonates with no adverse events such as overt bleeding noted.

**Table 1.**
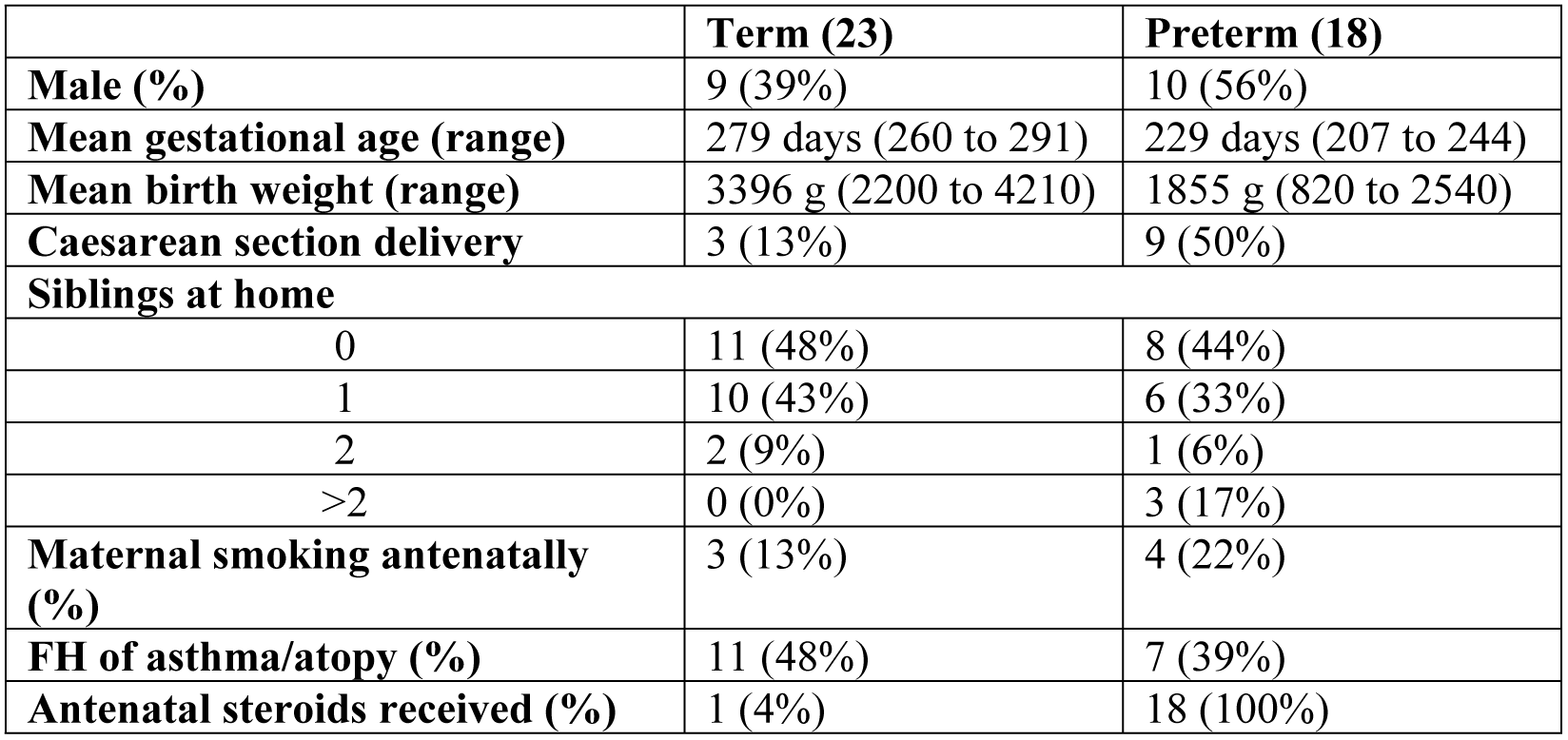
Perinatal and delivery characteristics of enrolled subjects. FH = Family history.

|  | **Term (23)** | **Preterm (18)** |
| --- | --- | --- |
| **Male (%)** | 9 (39%) | 10 (56%) |
| **Mean gestational age (range)** | 279 days (260 to 291) | 229 days (207 to 244) |
| **Mean birth weight (range)** | 3396 g (2200 to 4210) | 1855 g (820 to 2540) |
| **Caesarean section delivery** | 3 (13%) | 9 (50%) |
| **Siblings at home** |  |  |
| 0 | 11 (48%) | 8 (44%) |
| 1 | 10 (43%) | 6 (33%) |
| 2 | 2 (9%) | 1 (6%) |
| >2 | 0 (0%) | 3 (17%) |
| **Maternal smoking antenatally (%)** | 3 (13%) | 4 (22%) |
| **FH of asthma/atopy (%)** | 11 (48%) | 7 (39%) |
| **Antenatal steroids received (%)** | 1 (4%) | 18 (100%) |

Nasal AECs from infants at one-year old were collected with the infant held in a parent’s arms. A cepillo cell sampler brush (Deltlab SLU, Barcelona, Spain) was introduced into each nostril in turn using the same technique as described above. Repeat sampling was achieved for 24 infants (13 term, 11 preterm) at a median age of 12.5 months (IQR: 12-14.75 months, range: 12-22 months) (table 2). Follow-up was not achieved in 17 infants either because parents declined re-attendance or did not respond to re-attendance invitation. As for sampling at birth, the nasal brushing procedure was well tolerated by all infants with no adverse events.

**Table 2.** Clinical history of infants returning for repeat sampling after first year of life. FT = Full term, PM = Preterm, GA = gestational age, M= Months, RSV = Respiratory Syncytial Virus, URTI= Upper respiratory tract infection, IC= inhaled corticosteroids.

| Subject | Age (M) | Sex | Recurrent URTIs | Bronchiolitis | Atopy | Medications |
| --- | --- | --- | --- | --- | --- | --- |
| FT | 16 | F | Y | N | N | None |
| FT | 18 | F | N | N | N | None |
| FT | 22 | F | Y | N | Y | IC, β2 agonist |
| FT | 13 | F | Y | N | N | None |
| FT | 15 | M | N | N | N | None |
| FT | 15 | M | Y | Y (RSV) | Y | None |
| FT | 13 | F | N | N | N | None |
| FT | 15 | F | N | N | N | None |
| FT | 13 | F | N | N | N | None |
| FT | 14 | M | N | N | Y | None |
| FT | 12 | F | N | N | Y | None |
| FT | 14 | F | N | N | Y | None |
| FT | 12 | M | N | N | Y | None |
| PM (33wks, GA) | 12 | F | Y | Y (RSV) | N | None |
| PM (29wks, GA) | 12 | M | N | N | N | None |
| PM (33wks, GA) | 12 | M | N | N | N | None |
| PM (33wks, GA) | 12 | M | N | N | N | None |
| PM (34 wks, GA) | 13 | F | N | N | N | None |
| PM (29 wks, GA) | 12 | M | N | N | N | H2 antagonist |
| PM (32 wks, GA) | 12 | M | Y | N | N | None |
| PM (33 wks, GA) | 12 | F | N | N | Y | None |
| PM (33 wks, GA) | 12 | F | N | N | Y | None |
| PM (33 wks, GA) | 12 | F | N | N | Y | None |
| PM (33 wks, GA) | 12 | F | N | Y | Y | None |

### Processing of nasal samples and creation of WD-PNEC cultures

Each brush, with attached cells, was placed in sterile phosphate buffered saline (PBS) mixed with transport medium (DMEM, 0.1% Penicillin/streptomycin (1:1 V/V)). Paediatric nasal cells were removed from the brushes and expanded in airway epithelial cell basal medium supplemented with an airway epithelial cell growth medium supplement pack (Promocell, Heidelberg, Germany) using established methods.(5,18) Upon reaching 70-80% confluency, the cells were seeded at passage 3 onto collagen-coated 6 mm diameter Transwells (Corning, Tewksbury, Massachusetts) at a density of 1×10^5^ cells/Transwell. After reaching confluency, air-liquid interface (ALI) conditions were established and maintained until complete differentiation occurred. Following complete differentiation, which was defined by extensive cilia coverage and mucus production under light microscopy, transepithelial electrical resistances (TEER) were measured, as described previously.(5)

### Immunofluorescence microscopy for WD-PNEC characterisation

For WD-PNEC cultures, representative Transwells were fixed in 4% (v/v) paraformaldehyde (PFA) for 30 min at room temperature (RT). Fixed Transwells were stored in 70% ethanol at +4°C until used. To prepare for immunofluorescence staining, ethanol was removed and cells rinsed twice with 200-300 μL PBS added to the apical surface (pH 7.4). Cells were permeabilised using 0.2% (v/v) Triton X-100 (Sigma-Aldrich, ST. Louis, Missouri) in PBS for 1 h at RT and subsequently blocked with 0.4% (w/v) bovine serum albumin (BSA) (Sigma-Aldrich, ST. Louis, Missouri) in PBS for 1 h at RT. Cultures were next stained for β-tubulin, Muc5Ac and nuclei (DAPI). Briefly, 200 μL rabbit anti-Muc5Ac antibody (ab78660, Abcam, Cambridge, UK) (1:100 dilution in 0.4% (w/v) BSA (Sigma-Aldrich, ST. Louis, Missouri) in PBS) was added and incubated overnight at 4°C. Cultures were washed 3 times with 200-300μL PBS added to the apical surface (pH 7.4) for 5 min at RT. Two hundred μL anti-rabbit secondary antibody (A11011, Alexa-Fluor 568, Invitrogen, Waltham, Massachusetts) was added (1:500 dilution in 0.4% (w/v) BSA (Sigma-Aldrich, ST. Louis, Missouri) in PBS) and incubated at 37°C for 1 h. Cultures were further washed 3 times with 200-300μL PBS added to the apical surface (pH 7.4), and 200 μL Cy3-conjugated mouse anti-β-tubulin antibody (ab11309, Abcam, Cambridge, UK) added (1:200 dilution in 0.4% (w/v) BSA (Sigma-Aldrich, ST. Louis, Missouri) in PBS) and incubated at 37°C for 1 h. Following further 3 × 200-300μL PBS (pH 7.4) washes added to the apical surface, nuclei were stained using DAPI-mounting medium (Vectashield, Vector Laboratories, Burlingame, California). Fluorescence was detected by confocal laser microscopy (TCS SP5 Leica, Germany) or by UV microscopy (Nikon Eclipse 90i, Nikon, Surrey, UK).

Cells from one representative Transwell for each culture were trypsinised and the cell suspension was either smeared onto two microscope slides or 200-250 μL of cell suspension was added to a cytofunnel (EZ single cytofunnel, Thermo Fisher Scientific, Waltham, Massachusetts) and spun at 100 rpm (Using Thermo Shandon Cytospin 4 Cytocentrifuge) for 4 min onto a microscope slide. Smeared and cytospun slides were then fixed in 4% (v/v) PFA for 15 min at RT. Fixed slides were stored in the dark at −20°C until immunofluorescence was performed. Slides were stained either for ciliated cells (anti-β-tubulin) or goblet cells (anti-Muc5Ac). In brief, cells were permeabilised using 0.2% (v/v) Triton X-100 (Sigma-Aldrich, ST. Louis, Missouri) in PBS (pH 7.4) for 30 min at RT and subsequently blocked with 0.4% (w/v) bovine serum albumin (BSA) (Sigma-Aldrich, ST. Louis, Missouri) in PBS (pH 7.4) for 30 min at RT. One slide was stained for Muc5Ac by addition of 100 μL rabbit anti-Muc5Ac antibody (ab78660, abcam) (1:100 dilution in 0.4% (w/v) BSA (Sigma-Aldrich, ST. Louis, Missouri) in PBS) incubated overnight at 4°C, followed by 3 x 200 μL PBS (pH 7.4) washes for 5 min at RT before addition of 100 μL anti-rabbit secondary antibody (A11011, Alexa-Fluor 568, Invitrogen, Waltham, Massachusetts) (1:500 dilution in 0.4% (w/v) BSA (Sigma-Aldrich, ST. Louis, Missouri) in PBS) at 37°C for 1 h. A second slide was stained for β-tubulin by addition of 100 μL Cy3-conjugated mouse anti-β-tubulin antibody (ab11309, Abcam, Cambridge, UK) (1:200 dilution in 0.4% (w/v) BSA (Sigma-Aldrich, ST. Louis, Missouri) in PBS) incubated at 37°C for 1 h. Both slides next underwent 3 x 200 μL PBS (pH 7.4) washes and nuclei were stained using DAPI-mounting medium (Vectashield, Vector Laboratories, Burlingame, California). Quantification of ciliated, goblet and total DAPI+ cell numbers was undertaken for each slide by counting under fluorescent microscopy (Nikon Eclipse 90i; Nikon, Surrey, UK) and the proportion of ciliated and goblet cells relative to total DAPI+ cell numbers was determined.

### Freezing and defrosting of harvested nasal epithelial cells

Expanded paediatric nasal cells were trypsinised at passage 3, added to low glucose DMEM containing 5% (v/v) foetal bovine serum (FBS) and centrifuged for 5 min at 129 × *g*. Following supernatant removal, the resulting cell pellet was re-suspended in monolayer medium (epithelial cell basal growth medium, Promocell, Heidelberg, Germany) containing 10% FBS and 10% dimethylsulfoxide (DMSO) (Sigma-Aldrich, ST. Louis, Missouri) to give a final concentration of 1 × 10^6^ cells/mL. One mL aliquots of the cell suspension in cryovials were placed into an isopropanol cell freezing apparatus (Mr Frosty, Nalgene, Thermo Fisher Scientific, Waltham, Massachusetts) at RT and transferred to −80°C for 24 h, before being stored long-term in the gaseous phase of a liquid Nitrogen (N_2_) tank.

To resuscitate frozen primary nasal epithelial cells, vials were removed from the liquid N_2_, defrosted rapidly in a water bath at 37°C and the contents centrifuged at 129 × *g* for 5 min. The resulting cell pellet was re-suspended in monolayer medium (epithelial cell basal growth medium, Promocell, Heidelberg, Germany), transferred to a collagen-coated 75 cm^2^ flask (Thermo Fisher Scientific, Waltham, Massachusetts) and incubated at 37°C in 5% CO_2_. The generation of WD-PNEC cultures then proceeded as described above.

### Statistical analysis

Data are presented as means ± standard deviation and median, interquartile range (IQR) and range for skewed data. Statistical analysis was performed by a student’s paired or unpaired t-test unless otherwise stated. Statistical significance was set at a p-value less than 0.05 (* p < 0.05 or **p <0.01). Data were analyzed using GraphPad^®^ Prism 5.0 (GraphPad Software, Inc., La Jolla, CA).

### Ethics statement

Written informed consent was obtained from parents at recruitment. Study was approved by the Office for Research Ethics Committee Northern Ireland (ORECNI), (REC reference 14/NI/0056).

## Results

### WD-PNEC cultures from preterm and term newborn infants demonstrate similar differentiation schedules and success rates

Primary monolayers were successfully established in 18 term and 15 preterm infants (80%). One sample failed to grow due to fungal contamination during culture. For the remaining samples, culture failure was due to insufficient cells harvested following brushing. Following establishment of primary monolayer culture, the subsequent success rate of differentiation into WD-PNECs was 30/33 (91%). Of the three cultures failing to achieve differentiation no obvious cause was evident, but it may be due to low sample yield resulting in cell division beyond the upper limit at which successful cell differentiation can occur. Although numbers were low, there was no correlation evident between maternal antenatal smoking and cell culture failure. Overall, the rate of success for complete differentiation from initial sample acquisition was 15/23 (65%) for recruited term neonates and 13/18 (72%) for recruited preterm neonates. WD-PNEC differentiation was established by the formation of extensive cilial coverage in all quadrants of each Transwell with clear mucous production and no holes evident, as observed under light microscopy. No significant difference was noted in the median duration of time required to achieve differentiation in term (77.5 days; IQR: 74 to 94, range: 64 to 97 days) and preterm (80 days; IQR: 70 to 88, range 61 to 133 days) cultures.

### Term- and preterm-derived newborn WD-PNEC cultures are morphologically indistinguishable

Newborn term and preterm WD-PNEC cultures were indistinguishable under phase-contrast light microscopy (figure 1A). Fluorescent microscopy of fixed and stained cultures confirmed the formation of multi-layered pseudostratified cultures containing ciliated and goblet cells (figure 1B). The mean proportions of ciliated cells in fixed and stained Transwells were similar in term- and preterm-derived WD-PNEC cultures (29.2% and 34.2%, respectively), as were the goblet cell proportions (26.7% and 23%, respectively) (figure 1C). These proportions were comparable to mean proportions of ciliated and goblet cells observed in representative trypsinised Transwell smears and cytospins (figure 1D). Tight junction integrity, as demonstrated by robust transepithelial electrical resistance (TEERs) of ≥300 Ω.cm^-2^, was evident for most Transwells in both term and preterm derived WD-PNEC cultures, with no significant difference between cohorts (figure 1E). Three term WD-PNEC cultures had TEERs ≤300 Ω.cm^-2^; however these cultures demonstrated extensive apical cilia coverage and obvious mucous production with no holes evident under phase-contrast light microscopy.

**Figure 1.**
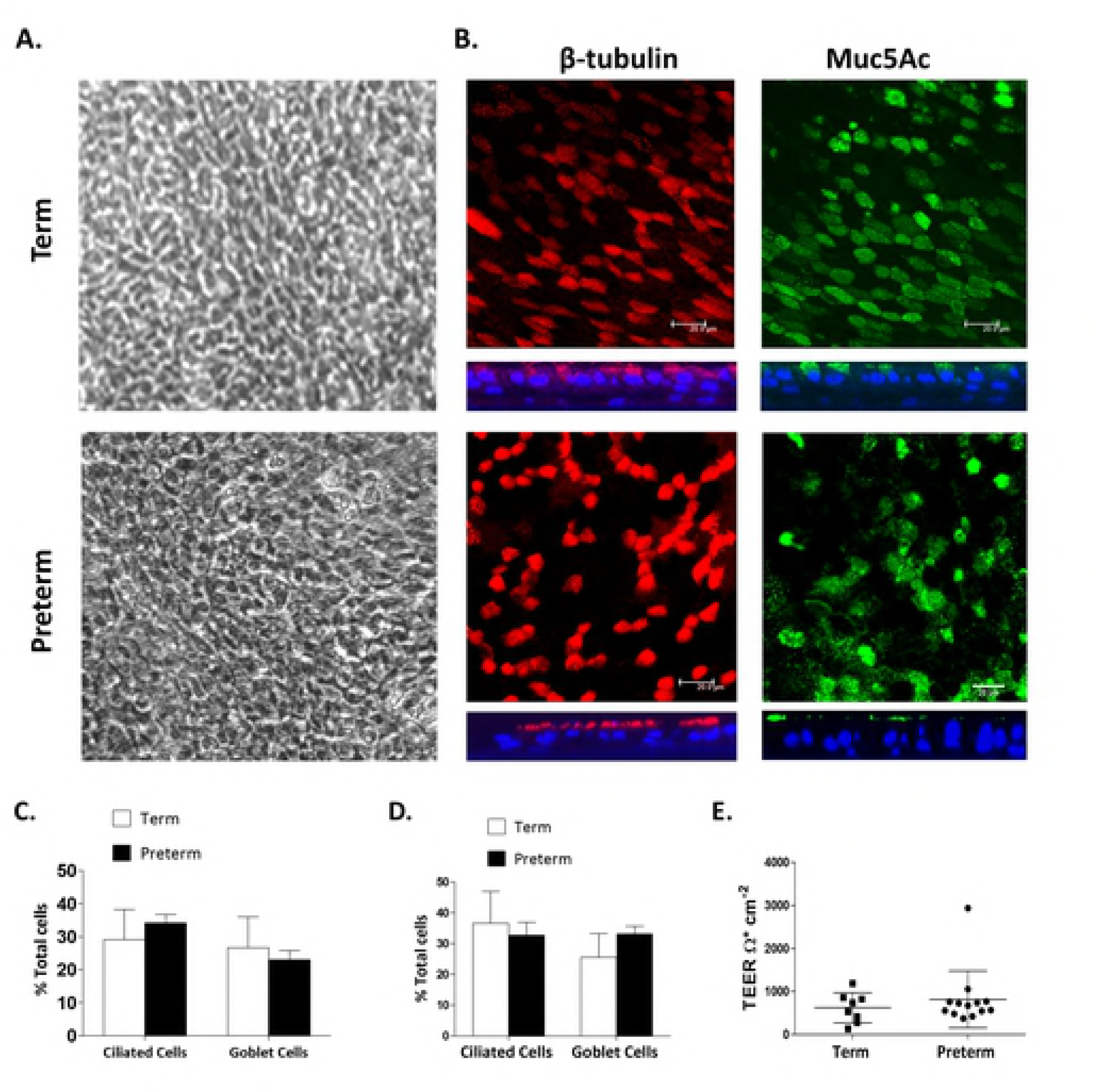
Morphology and differentiation status of newborn term and preterm WD-PNEC cultures. Cultures were monitored by (A) phase-contrast microscopy (magnification x20) or (B) confocal microscopy after staining for β-tubulin (ciliated cell marker) (red), Muc5Ac (goblet cell marker) (green), or nuclei (DAPI) (blue). For (B), square panels represent *en face* images, whereas rectangular panels represent orthogonal sections, with the apical side at the top (magnification x63, with x1.5 digital zoom, bar = 20 μm). (C) Transwell cultures from term and preterm donors (n=4 each) were fixed and stained for ciliated or goblet cells. Images from 5 fields/Transwell were taken at magnification x60 and ciliated, goblet cell and total DAPI^+^ cell numbers were counted using fluorescent microscopy. The % ciliated and goblet cells were determined relative to total DAPI^+^ cell numbers. Data are presented as mean (±SD). (D) Transwells cultures were trypsinized, contents fixed onto slides by smearing or use of cytospin funnels, as described, and stained for ciliated (anti–β-tubulin) and goblet (anti-Muc5Ac) and total DAPI^+^ cells (n=9 term, n=10 preterm). Ciliated and goblet cell numbers were expressed as a percentage of the total DAPI^+^ cell numbers. Data presented as mean (±SD). (E) Transepithelial electrical resistances (TEER) of neonatal term- and preterm-derived WD-PNECs. Measured by EVOM epithelial voltometer and presented as mean (±SD) Ohm.cm^-2^

### Nasal AECs from one-year old infants achieve complete differentiation faster than those from newborn infants

Primary monolayers were successfully established in 10/13 (77%) term and 9/11 (82%) preterm of repeat nasal brushing samples. Following establishment of primary monolayer culture the subsequent success rate of differentiation into WD-PNECs was 17/19 (89%). Of the two cultures failing to achieve differentiation no obvious cause was evident. Overall, the rate of success for differentiation from initial sample acquisition was 9/13 (69%) for one-year term infants and 8/11 (73%) for one-year preterm infants. Importantly, the time to achieve complete culture differentiation was significantly shorter for one-year cohort samples (median 63.5 days; IQR 49 to 72 days) than for birth cohort samples (median 80 days; IQR 72 to 133 days), p=0.0001 (2-tailed Mann-Whitney U).

### WD-PNEC cultures derived from one-year old infants demonstrate significantly reduced goblet cell content compared to their paired newborn-derived WD-PNECs

One-year WD-PNEC cultures were indistinguishable from newborn-derived cultures under phase-contrast light microscopy. As for newborn-derived WD-PNECs, there was no difference observed between the proportions of ciliated and goblet cells in term and preterm-derived cultures at one year (figure 2). However, we observed a significant decrease in mean goblet cell proportions at one-year compared to newborn-derived WD-PNECs for both term (11.7% vs 25.5%; p=0.0003) and preterm cohorts (8.9% vs 33.1%; (p<0.0001) when trypsinised Transwell cell smear proportions were examined (figure 2D).

**Figure 2.**
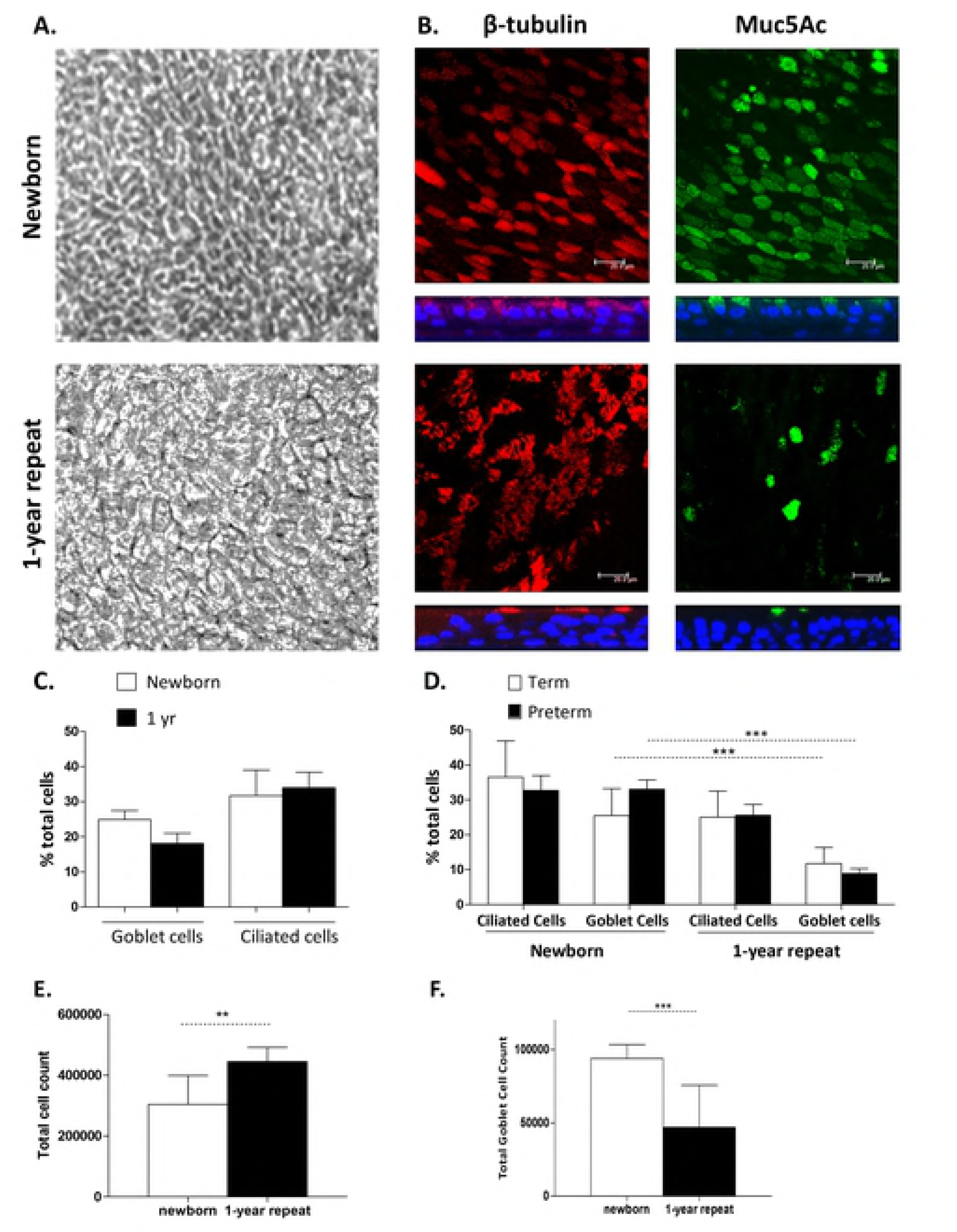
Morphology and differentiation status of birth and one-year repeat WD-PNEC cultures. Cultures were monitored by (A) phase-contrast microscopy (magnification x20) or (B) confocal microscopy after staining for β-tubulin (ciliated cell marker) (red), Muc5Ac (goblet cell marker) (green), or nuclei (DAPI) (blue). For (B), square panels represent *en face* images, whereas rectangular panels represent orthogonal sections, with the apical side at the top (magnification x63, with x1.5 digital zoom, bar = 20 μm). (C) Transwell cultures from birth and 1-year donors (n=8 each) were fixed and stained for ciliated, goblet and total DAPI^+^ cells. Images from 5 fields/Transwell were taken at x60 magnification and ciliated, goblet and total DAPI^+^ cell numbers were counted using fluorescent microscopy. The % ciliated and goblet cells were determined relative to total DAPI^+^ cell numbers. (D) Representative preterm and term Transwell cultures (newborn term n=9, preterm n=10, one-year term n=9, preterm n=6 donors) were trypsinised, contents fixed onto slides by smearing or use of cytospin funnels, as described, and stained for ciliated (anti–β-tubulin), goblet (anti-Muc5Ac), and total DAPI^+^ cells. Ciliated and goblet cell numbers were expressed as mean (±SD) % proportion of total DAPI^+^ cell numbers for each cohort. *** unpaired t-test, p<0.0001. (E) Transwell cultures were trypsinized and total cell count performed by trypan blue exclusion using a haemocytometer. Total cell numbers are presented as mean (±SD) for newborn (n=17) and 1-year cohorts (n=15), p= 0.0087 (unpaired t-test of difference in means). (F) Combined total goblet cell numbers were determined for trypsinised representative Transwell cultures. Data plotted for newborn (n=17) and 1-year (n=15) cohorts as mean (±SD) for each culture cohort. ***unpaired t-test of difference in means, p=0.0005.

We noted the total number of cells per Transwell culture was higher for the one-year versus newborn cohort (figure 2E). Therefore, to determine if the observed differences in goblet cell proportions could be explained by differences in total cell numbers within pseudostratified cultures we re-analysed the data as total goblet cell numbers for each donor. Consistent with the proportion data, combined total goblet cell count for all newborn-derived WD-PNECs (mean= 93,797 cells; SD 39427) was double that of one-year derived-WD-PNECs (mean= 46,969; SD 28607: t-test of difference in means, p=0.0014) (figure 2F). These data suggest the observed decrease in goblet cell proportions in one-year-derived WD-PNECs is not simply the result of increased total WD-PNEC culture cell numbers in the one-year cohort.

### Nasal primary AECs successfully differentiate after storage in liquid nitrogen and are morphologically indistinguishable from freshly differentiated AECs

We next sought to determine if nasal AECs could be successfully frozen and stored, as this vastly increases the versatility of this model for future research into the origins of airway disease. Newborn and one-year old nasal samples stored in liquid N_2_ for approximately one year were defrosted as described and expanded to enable Transwell seeding and differentiation.

Both samples successfully differentiated in a similar fashion, requiring one additional week to achieve complete differentiation compared to fresh counterparts. Differentiation was evidenced by the formation of extensive cilial coverage in all quadrants of each Transwell with clear mucous production under light microscopy (figure 3A). The extra week was due to additional time needed to enable cell expansion in monolayers prior to Transwell seeding.

**Figure 3.**
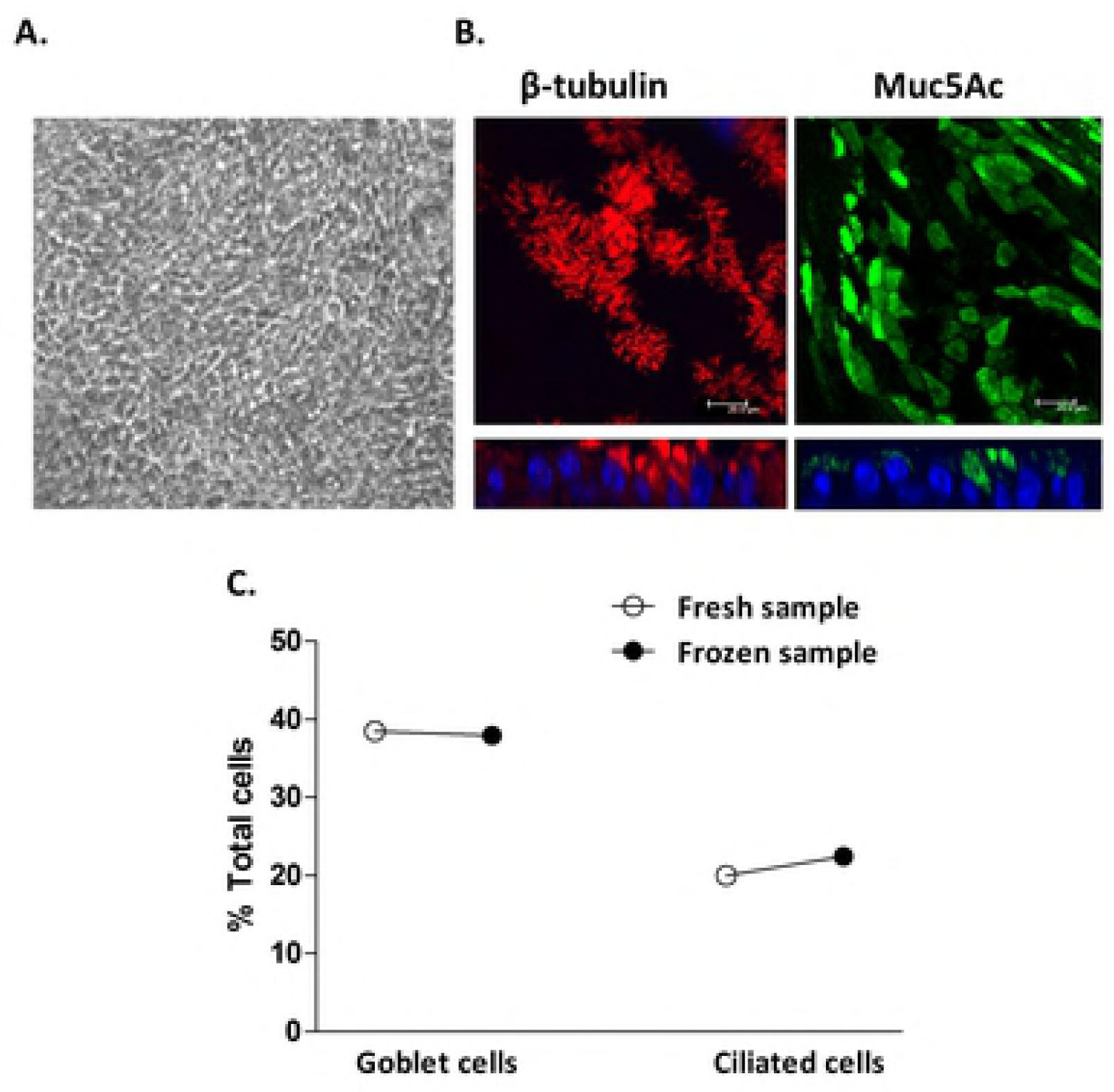
Morphology and differentiation status of WD-PNEC cultures derived from epithelial cells frozen at passage 3. Cultures were visualised by (A) phase-contrast microscopy (magnification x20) or (B) confocal microscopy after staining for β-tubulin (ciliated cell marker) (red), Muc5Ac (goblet cell marker) (green), or nuclei (DAPI) (blue). For (B), square panels represent *en face* images, whereas rectangular panels represent orthogonal sections, with the apical side at the top (magnification x63, with x1.5 digital zoom, bar = 20 μm). (C) Representative Transwell cultures derived from n= 1 newborn donor either differentiated “fresh” or following a period of storage in liquid nitrogen “frozen” were fixed and stained for ciliated, goblet and total DAPI+ cells. Images from 5 fields/Transwell were taken at x60 magnification and ciliated, goblet and total DAPI^+^ cell numbers were counted using fluorescent microscopy. The % ciliated and goblet cells were determined relative to total DAPI^+^ cell numbers.

Transepithelial electrical resistances (TEERs) >300 Ohm.cm^-2^ were recorded for Transwells in both cultures indicating robust epithelial cell tight junction formation. As described above for freshly processed nasal epithelial cells, fluorescent microscopy of fixed and stained WD-PNEC cultures derived from frozen cells confirmed extensive ciliated and goblet cell contents (figure 3B). Importantly, proportions of goblet and ciliated cells in frozen samples were very similar to those from the same donor cultured without freezing, with the unusually high goblet cell content from this donor being reproduced (figure 3C). While this was limited to cells from a single donor, our data demonstrate a proof of principle for the use of freezing newborn nasal epithelial cells for subsequent culture, differentiation and experimentation.

## Discussion

Very little is known regarding the development of the human airway epithelium in early life and how innate immune responses to airborne pathogens and allergens change with increasing age. We are not aware of any previously published studies describing the successful differentiation and characterisation of human newborn AECs and as such this work represents a unique model for studying early life airway epithelium immune responses.

As described by Miller *et al*, we found the nasal brushing procedure to be safe and well tolerated by infants. Our culture success rate for the newborn cohort of 80% for monolayer cultures is similar to passage 3 success rates for Miller *et al.* (7) Sample success rates for one-year old infants were almost identical to rates of success for the birth cohort, indicating consistency of sampling and culture technique throughout the study. The generation of morphologically authentic WD-PNECs from newborn AEC monolayer cultures that demonstrated typical pseudostratified columnar epithelium, goblet and ciliated cell generation, mucous production and robust TEERs was achieved with a high success rate (91%). This indicates our technique is reliable in enabling establishment of WD-PNECs from newborn samples. While the relatively small sample size in our current study limits correlation of findings with *in vivo* clinical characteristics, including relevance of atopy or susceptibility to recurrent upper respiratory tract infection, the consistency of this technique highlights its accessibility for use in future studies.

WD-PNECs generated from term and preterm newborn infants were morphologically and physiologically indistinguishable. Differentiation was achieved in a similar time frame for both. Importantly, this indicates no obvious impact of gestational age on AEC growth and differentiation. However, intriguingly, we detected morphological and differentiation differences between WD-PNECs generated from newborn and one-year old infants, suggesting age-related developmental changes occur in nasal AECs during the first year of life.

The difference in the speed at which full WD-PNEC differentiation is achieved between newborn and one-year samples is an interesting finding. Indeed, it may represent an increased capacity for nasal AEC proliferation and differentiation at one-year compared to the neonatal period. Differences in epithelial cell proliferation speed and functionality between neonatal, later postnatal age and adulthood have been observed in other epithelial tissues, namely, in human skin keratinoctyes and in mice gastrointestinal epithelium.(19,20) Mice studies have demonstrated that intestinal epithelial cell turnover is much faster in older animals compared to neonates and it has been proposed this difference is regulated intrinsically by genetic programming, mediated in part by the transcriptional repressor Blymphocyte-induced maturation protein 1 (Blimp1).(21) It is possible that similar intrinsic programming determines the rate of human AEC proliferation and differentiation. However, we do note that nasal sampling in newborn infants is limited by their comparatively smaller nasal passages, which in principle could result in reduced initial AEC basal cell yield. This might require more cell doublings and therefore longer times to achieve sufficient cell numbers for differentiation. We did not directly address basal cell numbers in brushing as it would necessitate sacrificing the sample. Instead, we observed cell adherence to the collagen-coated plastic from the brushes and found no obvious differences between brush samples from newborns and one-year olds.

In regards to proportions of ciliated and goblet cells, we observed that WD-PNECs derived from the one-year cohort in this study were similar to previous work in WD-PNECs from older infants.(6) In contrast, we observed significantly higher proportions of goblet cells in the newborn-derived WD-PNECs compared to one-year WD-PNECs. This is a valuable finding that has not previously been described. We are aware of only one study examining the proportion of mucous secreting cells in human newborn airway epithelium performed on post-mortem paraffin-embedded lung sections. This work only included three infants over 6 months old, thus its comparability with our work is limited.(22) Respiratory viral infections, such as RSV, result in excess mucous production and formation of thick mucous plugs, which contribute to the pathogenesis of respiratory viral diseases.(23) Thus, our finding of increased goblet cell content in newborn airways compared to older infants could contribute to increased mucous plug formation during viral infections and may explain, in part, the increased frequency of severe respiratory viral disease in very young infants. Indeed, there is some evidence to support such a hypothesis in the neonatal mouse model, where increased levels of Muc5Ac expression detected at baseline in neonatal versus adult mice was shown to further increase in response to human rhinovirus infection.(24,25) Increased goblet cell numbers are also observed in asthmatic airways and it is theorised overexpression is due to release of T helper 2 (Th2) cytokines in response to allergen-induced inflammation.(26) Interestingly, during fetal development and the early newborn period, human innate immune responses are biased towards Th2-cell polarising responses.(27) It is possible Th2 bias may offer an explanation as to why newborn WD-PNECs demonstrate increased goblet cell proportions. Further work is needed to determine the mechanisms contributing to this increase which may yield insights into diseases, such as asthma, where goblet cell hyperplasia is a feature.

Due to the small numbers involved in this study we could not determine any significant correlation between parental reports of severe or recurrent URTIs and goblet cell proportions. Furthermore, we recognise that parental reporting of clinical symptoms is limited by potential recall bias which must be considered when interpreting our clinical characteristics data.

One potential limitation with our model is the use of nasal rather than bronchial AECs and there is debate regarding the validity of nasal AECs as surrogates to investigate lung disease.(28) Some studies of airway inflammation in adult populations have not demonstrated that nasal AECs can substitute for bronchial AECs, such as in COPD.(29) However, studies of bronchial AECs in children have thus far been largely opportunistic, sampling children when anaesthetised to allow access to the bronchial epithelium. (8,30,31) Evidently, as we have discussed earlier, bronchial sampling is not appropriate in newborn infants who are otherwise well. Moreover, use of bronchial AECs would not permit repeated sampling within the same healthy subject, which is one of the most attractive exploitations of our model. Furthermore, indistinguishable epithelial morphology has been reported in paired nasal and bronchial AEC cultures from both adults and children, while more recent work in paediatric asthma, atopy and respiratory syncytial virus infection demonstrated that nasal AECs act as reasonable surrogates for lower airway AECs.(6, 32–34)

In conclusion, we present the first description of morphologically and physiologically authentic WD-PNEC cultures generated from term and preterm newborn infants. We demonstrated this to be a safe, minimally invasive method that can be performed consistently with high rates of success. Attaining follow-up in over half of initially enrolled infants provided sufficient numbers to enable robust comparisons between groups and demonstrates the feasibility of sequential sampling of subjects for future research studies. Accordingly, our model of the neonatal airway presents a unique opportunity to study differences contributing to the increased severity of respiratory infections seen in this age group compared to older infants. Furthermore, the successful protocol for freezing nasal AECs to enable differentiation at a later date presents an opportunity for even greater flexibility in using these samples for research into childhood respiratory illnesses. For instance, this methodology offers a unique opportunity to store ‘‘naıve’’ AECs and differentiation may be performed at a later date once absence/presence of subsequent respiratory disease, e.g. asthma, has been established. Thus, newborn WD-PNECs represent a unique opportunity to study innate immune responses of human airway epithelium from birth, and have significant and exciting potential applications in elucidating the early origins of respiratory disease.

## Acknowledgements

We are extremely grateful to the parents and infants whose participation in this study made it possible. We would also like to thank Isobel Douglas, research nurse, for assistance in training in the technique of nasal brushing sampling.

## Notes

**Statement of financial support:** This works was supported by Wellcome Trust grant number 104516/Z/14/Z to Helen Groves.

